# Automatic surveillance of *Escherichia coli* bacteriological strains within clinical settings

**DOI:** 10.1101/2024.11.05.622049

**Authors:** Rafael Rodríguez Palomo, Alejandro Guerrero-López, Carlos Sevilla-Salcedo, David Rodríguez-Temporal, Belén Rodríguez-Sánchez, Vanessa Gómez-Verdejo

**Affiliations:** Departamento de Teoría de Señal y Comunicaciones, Universidad Carlos III de Madrid, Madrid, España; Instituto de Investigación Sanitaria Gregorio Marañón, Madrid, Spain

**Keywords:** Surveillance, Escherichia coli, Automatic, Machine Learning, Clinical microbiology

## Abstract

Healthcare-associated infections (HAIs) are a significant concern within hospital environments, with the World Health Organization (WHO) identifying them as a major source of bacteriological infections. HAIs affect millions of patients annually, leading to substantial morbidity and mortality. However, a significant proportion of HAIs are preventable through early detection and appropriate intervention and isolation. Traditional methods for identifying bacterial species and strains, such as antigen tests, are often time-consuming and hamper the realtime tracking of outbreaks. The matrix-assisted laser desorption ionization-time of flight mass spectrometry (MALDI-TOF MS) technique has emerged as a more rapid and precise alternative, though it remains limited by manual identification processes and database constraints. Recent advancements have demonstrated the potential of combining MALDI-TOF MS data with machine learning (ML) to enhance in silico identification speed. In this paper we propose the first method for unsupervised *Escherichia coli* novel strain detection through the application of a efficient PIKE tailored to strain identification, spectral clustering techniques and a one-class support vector machine (OCSVM) novelty detection model, presenting promising results.

## Introduction

According to the World Health Organization (WHO) hospital environments are one of the most powerful sources of bacteriological infections (1), also referred as nosocomial infections or healthcare-associated infections (HAIs). HAIs are defined as infections acquired by patients during their stay in a hospital. Although some of these infections are simple to treat, others might cause complications to the preexisting conditions of the patients.

It is estimated that more than 3.5 million cases of HAIs will occur in the European Union (EU) every year, resulting in deaths and 2.5 million disability adjusted life years (2). Moreover, the widespread and uncontrolled use of antimicrobial drugs within hospital environments has caused a spike in the amount of antibiotic-resistant bacteria, with up to 71% of HAIs resulting in this type of infection (2). However, 50% to 70% of these nosocomial infections are estimated to be avoidable (1, 2). Hence, the ability to early detect the bacteriological species and, specially, the strains present in a patient at the moment of admission is crucial to allow for the proper measures to be taken, such as isolating the patient to avoid further outbreaks or helping clinicians to select specific antibiotics during treatment.

In standard clinical practice there are several methods for performing bacteriological species and strain identification such as: direct morphological observation under a microscope, antigen test or the culture method (3). However, there is a method, used as standard clinical practice during patient admission, that provides faster and more accurate results in combination with overnight culture methods, the matrix-assisted laser desorption ionization-time of flight mass spectrometry (MALDI-TOF MS) technique (3).

MALDI-TOF produces a proteomic spectrum with enough information to identify certain bacteriological species by means of bacteria databases. However, the identification is performed manually by trained professionals relying on a small number of features, such as peak position and height, to identify species that have been empirically related with those attributes (4). Moreover, MALDI-TOF MS based identification is hampered by the extent of the database at use and the high MALDI-TOF device and observerwise intervariability. Consequently, to confidently confirm whether an isolate belongs to one strain or another it is necessary to perform genetic sequencing of the sample.

MALDI-TOF data has been successfully used in the past in combination with machine learning (ML) techniques to tackle certain problems, such as antibiotic resistance prediction (5, 6), species identification (7) or bacteria ribotyping (8).

In this paper we aim to leverage the power of kernel methods, clustering and novelty detection techniques in combination with *Escherichia coli* (*E. coli*) MALDI-TOF MS to propose a pipeline for the automatic and unsupervised identification and detection of novel strains, as well as providing an automated method for the surveillance of *E. coli* outbreaks within clinical settings. In addition, we seek to identify the *E. coli* strains that exhibit a higher virulence.

The rest of the document is organized as follows. Section 2 presents the state-of-the-art related to the use of MALDI-TOF data in combination with artificial intelligence techniques and, specifically, the relevant work associated with the identification of bacteriological strains. Section 3 provides detailed descriptions of materials and methods used in the study. Section 4 contains the description of the obtained results. Finally, section 5 presents the discussion of the results while section 6 provides an overview of the conclusions.

## Related Work

Artificial Intelligence (AI) techniques have been successfully used in the past in combination with MALDI-TOF MS data to tackle certain problems in microbiology, namely antimicrobial drug resistance prediction and bacteria identification. In (4), C. Weis et al. provide a review of studies employing AI and ML techniques for the identification of bacteriological species and for antimicrobial susceptibility testing. They found 36 studies, 27 of them used ML for species identification while 9 of them focused on antimicrobial resistance prediction.

The following sections contain a collection of more recent studies not contained in the mentioned review as well as the most relevant studies for this project.

### A. Antibiotic resistance prediction

In (5), the authors propose a multi-view heterogeneous Bayesian model for the prediction of antibiotic resistance in *Klebsiella pneumoniae* isolates. E. Gato et al. (9) propose a dual pipeline for bacteria identification and antimicrobial drug prediction from MALDI-TOF MS data based on four models: partial least squares discriminant analysis (PLSDA), support vector machine (SVM) with and without a principal components analysis (PCA) applied, k-nearest neighbor (KNN) with and without a neighborhood components analysis (NCA) and a random forest (RF). C. Weis et al. (6) compare different classifiers, including a LightGBM, a Logistic Regressor (LR) and a Multi Layer Perceptron (MLP) to tackle resistance prediction on the Database of ResIstance against Antimicrobials with MALDI-TOF Mass Spectrometry (DRIAMS), containing over 300.000 mass spectra. In (10), the authors present the peak information kernel (PIKE), a similarity measure tailored to MALDI-TOF MS data. They combine the PIKE with Gaussian Process classifiers to predict antibiotic resistance while providing an estimation of the uncertainty, outperforming the pre-existing methods.

### B. Bacteria identification

The majority of the pre-existing literature focuses on the identification at the species level. In (11), the authors use a supervised learning approach to train a SVM with a Gaussian RBF kernel and an RF, reporting accuracy values ranging from 94% to 98% for the identification of *Leuconostoc* and *Fructobacillus* species. More recently, Y. Li et al. (12) propose a pipeline for *Listeria* species MALDI-TOF MS dimensionality reduction and feature extraction based on a denoising autoencoder (DAE) and a convolutional neural network (CNN). They test the feature extraction methods with several classifiers, where the combination of a DAE and a SVM as classifier yield the best results in every test metric. In (13), E. Kim et al. test the ability of a KNN combined with a PCA, a SVM with an RBF kernel and an artificial neural network to differentiate between the *cibaria* and *confusa* species of the genus *Weissella*, with the SVM outperforming the other methods.

However, to our knowledge, only the following studies focus on subspecies or strain identification. T. Mortier et al. (14) propose a large-scale benchmark of MALDI-TOF MS bacteria identification considering different cases, one of them in the novel strain case, where a set of unknown strains are introduced in the test set. They compare several models using a flat or a hierarchical classification reporting an accuracy of 84.78% using a local outlier factor (LOF) model, with no significant difference between the flat and hierarchical classification approaches. In (15), Rodríguez-Temporal et al. use supervised learning techniques in combination with labeled MALDI-TOF MS strain data to train a RF to identify different *Mycobacterium abscessus* subspecies (*abscessus, bolletii* and *massiliense*). They report that 96.5% of isolates were correctly identified and that geographical origin of the subspecies impact the spectra peak distribution.

Although there exists evidence that MALDI-TOF MS data can be used in combination with ML to identify different strains, there is a clear lack of studies focused on the sole task of the identification of novel strains in an unsupervised manner. This has the potential to facilitate the identification process without the need for previously labeled data.

In this article we aim to extend the capabilities of the base PIKE to adapt its implementation to strain identification and combine it with unsupervised clustering techniques and novelty detection methods to propose a pipeline for the rapid identification of novel and previously known bacteriological strains, as well as providing a tool for temporal detection of novel strains in clinical settings.

## Materials and Methods

### C. Database

We use a pre-existing and not yet public database from Hospital General Universitario Gregorio Marañón (HGUGM), acquired between 2018 and 2019. The dataset consists of 1525 preprocessed *E. coli* MALDI-TOF MS samples from a MALDI Biotyper (Bruker Daltonics) system. Each sample is associated with a series of labels, described in Table 1.

**Table 1.** Database Labels.

| Label | Description |
| --- | --- |
| Patient age | Numeric variable ranging from 0 to 102 |
| Patient sex | Categorical variable, male or female |
| Acquisition date | Date variable ranging from 2018-01-06 to 2019-12-08 |
| Department | Categorical variable representing the hospital department where the sample was acquired, variable has 111 levels |
| Sample type | Categorical variable representing the sample type (e.g. blood, urine, etc.), variable has 7 levels |
| Antibiotic resistance | Categorical variable classifying samples as resistant or non-resistant, resistant samples are subdivided in Extended Spectrum Beta-Lactamase (ESBL) resistance or Carbapenemase (CP) resistance |

All 1525 samples are previously preprocessed using the MALDIquant R library (16) following the standard pipeline for MALDI-TOF MS data. In addition, two extra steps are carried out in every sample: (i) MALDI-TOF MS m/z values are clipped to the range 2-20 kDa, so each MALDI contains 18000 data points and (ii) m/z values are approximated to its closest integer value, assigning the mean intensity value to each discrete m/z position.

There are no ground truth labels indicating the different *E. coli* strains present in the dataset. An objective of this study is to identify these strains in order to find the most virulent variants.

### D. Preprocessing

Before feeding the MALDI-TOF MS data to the kernel we apply several preprocessing methods to identify the best pipeline in terms of bacterial strain identification.

#### D.1. Binning

As presented in (6, 10), we use binning to reduce the dimensionality of the spectra and to speed up the computation of the kernel. Constant binning is the most widely used option in the literature, we follow the same approach.

After binning we normalize the scale of all spectra so that the sum of all intensities within a MALDI-TOF spectrum sums up to 1, maintaining the scale yielded by the MALDIquant pipeline.

#### D.2. Noise removal

MALDI data is sparse, in the sense that the majority of the m/z positions present low and almost-zero intensities while only some positions present relevant peaks. These almost-zero intensities pose a problem in the following scale normalization step, causing this step to yield very high intensity values in the relevant peaks positions to satisfy Eq. (1) in the low intensity ones. This problem can be mitigated by performing noise removal.

In this paper we propose that for each spectrum the mean intensity is calculated. Then, intensity values lower than the spectrum mean minus one standard deviation are flattened to zero. Although it might seem that flattening peaks is an aggressive approach it must be noted that MALDI data is sparse, resulting in very low mean values. Peaks are always above the mean, being unaffected by the noise removal step.

#### D.3. Scale normalization

This step is introduced by C. Weis et al. in (10) to ensure the successful calculation of the PIKE similarity measure. To avoid the product of two intensities becoming progressively smaller, all spectrum intensities must satisfy:

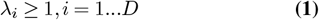

Where *λ*_*i*_ is the MALDI-TOF intensity at position or bin *i − th* and *D* is the length of the spectra.

### E. PIKE

We use the PIKE proposed by Weis et al. in (10) to leverage its OOD rejecting capabilities. The PIKE is specifically designed to work with sets of tuples, representing m/z positions and their corresponding intensities, without the need for feature vectors. Thus, it is able to manage similarities and interactions between different peaks in a MALDI-TOF spectrum. The PIKE is defined as:

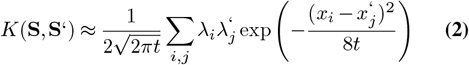

Where *S, S*’ are a pairs of MALDI-TOF spectra, *λ*_*i*_ and *λ*_*j*_ are two individual peak intensities of *S* and *S*’, respectively, and *x*_*i*_ and *x*_*j*_ are their respective location in the m/z axis or, as in our case, the bin axis.

The only adjustable parameter is *t*, the smoothing factor that controls the influence of other peaks within the spectrum. This parameter can be adjusted automatically by standard optimization techniques in supervised learning scenarios. For the purpose of this study we use a smoothing factor of 4, as suggested in (10).

### F. PIKE for strain identification

We introduce two new modifications to the previously described PIKE in order to adapt its implementation and formulation to the present problem of strain identification.

#### F.1. Efficient PIKE (E-PIKE)

The main limitation of the original PIKE implementation is its scalability, to compute the similarity measure between two spectra all pairs of peaks must be compared. To mitigate this problem the authors of the PIKE propose to use a topological filtering strategy to only consider the most relevant peaks. However, in the problem of bacteriological strain identification the most relevant peaks of the spectra only serve to identify the species level, while the smallest and, often, the less relevant peaks are the ones that differentiate one strain from another.

To overcome this issue we introduce E-PIKE, and efficient PIKE implementation that allows for the computation of the similarity measure using the whole spectrum. Assuming, as in our present case, that due to the binning process all spectra have the same length and that m/z values are identically distributed we can precompute the exponential term in Eq. (2), obtaining a universal distance array *d* that captures the distance between every pair of m/z points. Moreover, only those intensities belonging to m/z data points that are closer than a certain distance threshold *th* are considered for the computation of a similarity measure between two MALDI spectra. In the present study we opt for an arbitrary threshold *th* of 1*e*^*−*6^, which we have found to be an appropriate value according to the distance array *d* yielded by the exponential and the chosen smoothing factor.

#### F.2. Standard peak removal (SPR)

With the help of the Instituto de Investigación Sanitaria Gregorio Marañón (IiSGM) from HGUGM common peaks in every spectrum are identified, these peaks have been empirically proven to be present in almost every *E. coli* MALDI-TOF MS spectrum representing the species level and are often associated with ribosomic proteins. We hypothesize that removing these common peaks might help in the strain identification task, amplifying the subspecies information contained within these less relevant peaks.

In this regard we introduce Standard Peak Removal (SPR), a modification of the PIKE’s base method to remove common peaks in kernel space.

Let *X* be an array of shape (*N, D*) containing the *N E. coli* MALDI-TOF samples, where each row contains a preprocessed spectrum of length *D. K*_*SP R*_ is the resulting kernel without common peaks removed in kernel space, *K*_*noSP R*_ is the original PIKE computed between *X* and itself as per Eq. (2), *K* is the resulting (*N*, 1) kernel between *X* and *X*_*CP*_, an artificially made MALDI-TOF spectrum tailored to our database and, finally, *K*_*CP*_ is the resulting (1, 1) kernel between *X*_*CP*_ and itself.

To broadcast the different kernel dimensionalities we include the following support matrices: 1_*N*_, an all ones matrix with shape (1, *N*) and 1_*NN*_, an all ones matrix with shape (*N, N*) . Common peaks are removed in kernel space according to:

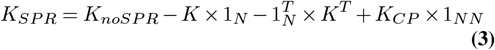

*X*_*CP*_ presents non-zero intensities only in the bins that contain common peaks. This intensities are assigned based on the median intensity of the ribosomic bins, defined by IiSGM, of the original data taking into account all 1525 available samples. The same preprocessing pipelines are applied to *X*_*CP*_ and the original MALDI-TOF data, except for the noise removal step that is only applied to the latter. Fig. 1 shows the representation of *X*_*CP*_.

**Fig 1.**
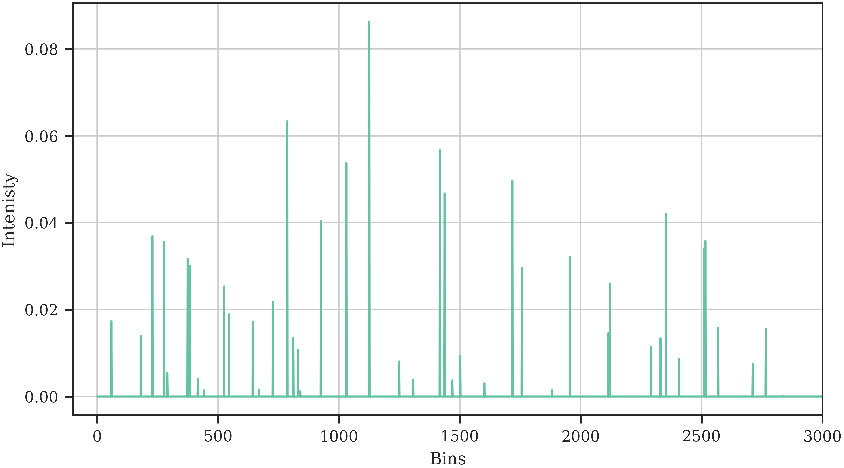
Representation of *X*_*CP*_, zoomed in the range [0, 3000]. There are no common peaks beyond bin 3000.

Moreover, to validate our SPR proposal we consider a third and simpler option. Instead of removing common peaks in kernel space we mask them before binning. To ensure that the common peaks are masked we consider a window of *±* 30 Da from the center of the peak. The same preprocessing consisting of binning, noise removal and scale normalization is applied to this pipeline. For this point onward we will refer to this third pipeline as the masked peaks MALDI-TOF pipeline.

Code implementing the E-PIKE and the SPR pipeline is publicly available in (17).

### G. Postprocessing

The resulting kernel is a similarity matrix. However, elements in the diagonal are not equal nor unitary. We employ a cosine similarity postprocessing step to transform our similarity matrix into a standard affinity matrix, ensuring that the diagonal elements are set to 1 and that the matrix value range is [0, 1], facilitating the interpretation of the results. Each element in the resulting kernel matrix is transformed according to:

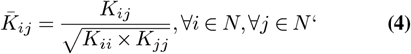

Where 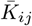 is the normalized kernel value at position [*i, j*], *N* and *N* ‘are the number of samples contained in the first and second MALDI-TOF arrays used in the kernel computation, respectively. The whole pipeline, from raw MALDI-TOF MS data to the kernel affinity matrix is summarized in Fig. 2.

**Fig 2.**
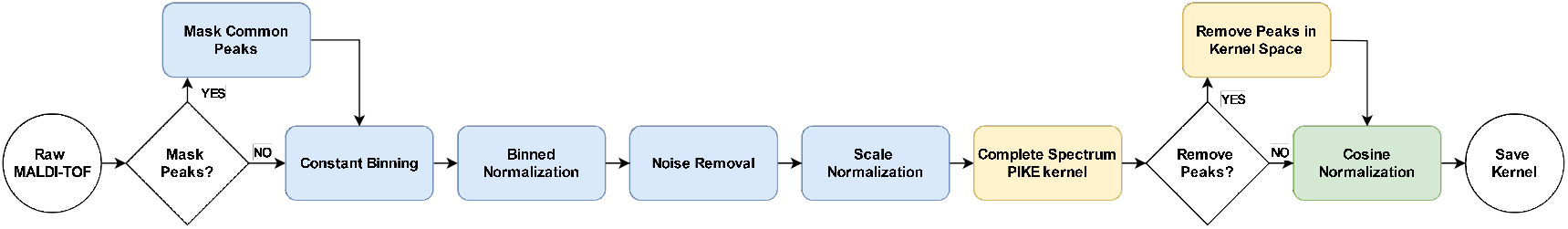
Pipeline from raw MALDI-TOF MS spectra to kernelized data.

### H. Spectral clustering

We use spectral clustering (18) in combination with the precomputed similarity matrices obtained from the E-PIKE to find groups of similar MALDI-TOF *E. coli* spectra. Spectral clustering, in combination with kernel methods, has been proven to be effective in a wide and varied range of applications, such as image segmentation (19–21), industrial maintenance prediction (22) or power grid modeling (23, 24).

Spectral clustering groups data by, firstly, embedding a Laplacian matrix into a low-dimensional space and, secondly, performing a traditional clustering technique (e.g. k-Means) of the components of the eigenvectors in the low-dimensional embedding space. It is particularly powerful for detecting clusters that have non-linear structures. The kernel formulation of spectral clustering has the advantage of being easily tailored to specific applications by the selection of an appropriate kernel. Moreover, it is able to accurately predict the group membership of unseen samples (23).

The number of clusters is selected according to the mean silhouette score. The silhouette score tests the goodness of a partition when no ground truth labels are available. It uses the mean intra-cluster distance and the mean nearest-cluster distance to evaluate the cohesion and separation of each sample, providing a quantitative estimation of the best number of groups in a dataset.

The silhouette score value is in the range [*−* 1, 1] and it is maximized when members within the same cluster are very similar among them and very dissimilar to samples in other groups.

### I. Temporal novelty detection

We use novelty detection to perform temporal detection of novel strains and outbreaks. Two novelty detection methods are tested: a One-Class SVM (OCSVM) and an Isolation Forest (iForest).

#### I.1. One-Class SVM

It is an unsupervised novelty detection model mainly used when only one class (the common class) is known. The OCSVM model detects outliers by learning a decision function that separates the common data from the rest of the space. There are two possible formulations of the OCSVM decision function (25): the separating hyperplane that finds the maximum margin hyperplane to separate the data from the origin and the separating hypersphere that tries to encapsulate all the training data within a sphere of the smallest possible radius.

The OCSVM model can be used, and it has been proven to be more effective (25), in combination with appropriate kernel methods to project data distributions into high dimensional feature spaces (26).

Thus, we propose to use the OCSVM model in combination with the PIKE and the previously computed spectral clustering for the novel strain detection task.

#### I.2. Isolation Forest

This model, introduced by Liu et al. in (27), proposes a new approach to novelty detection by isolating the anomalies instead of learning the common class. It builds an ensemble of trees that isolates observations by splitting the data based on its attributes (28). Samples that require, in average, a low number of splits to be isolated are considered anomalies. iForests have been successfully used in many diverse real-world applications in the past (28–32). This model poses several advantages to other anomaly detection methods: it is not a distance nor density-based model, being more computationally efficient than the alternative methods, it escalates well with dataset size and feature vector length and it leverages the power of ensemble methods to anomaly detection (27).

Although the classic formulation of the iForest model does not consider kernel compatibility, Li et al. (33) showed that combining kernel methods and iForests outperforms other state-of-the-art methods and that anomalies are easier to isolate from background in kernel space. Hence, following their approach, we combine the precomputed E-PIKE matrix with an iForest to compare its performance with that of the OCSVM.

### J. Experiments

Several experiments are carried out to test the capabilities, in terms of strain identification and detection, of our proposed modifications of the PIKE and preprocessing pipeline. We adhere the following process.

#### J.1. Preprocessing and clustering

First, we compute the E-PIKE taking into account all three possible peak removal scenarios:

- Without SPR.
- With SPR.
- With masked MALDI-TOF data.

We use 6000 bins of size 3 in the binning step. The results obtained from our proposed pipelines are then compared in terms of cohesion based on the silhouette score.

The resulting E-PIKE similarity matrices are clustered using the same number of clusters in every possible scenario, where the number of clusters is selected so that every pipeline yields a similar silhouette score. This will allow us to perform an in depth comparison of the clustering results as follows:

- First, we find which samples are always classified in the same cluster and which samples change from pipeline to pipeline.
- Second, we plot the mean MALDI-TOF spectrum of the common samples per cluster to identify trends in the spectra.
- Then, for each pipeline and cluster we plot the mean MALDI-TOF spectrum of the non-intersecting samples as assigned by each pipeline.

With these figures we will perform an in depth comparison to find what MALDI-TOF features are more relevant for the sample tipyfication in each pipeline, selecting the most reliable one.

Finally, with the selected pipeline is chosen we perform the novel strain detection analysis. In addition, all possible kernelization pipelines will be reviewed by IiSGM to confirm our results.

#### J.2. Novel strain identification

Here we compare the performance of the OCSVM and the iForest models in the task of novel strain identification, using the affinity matrix and cluster labels of the selected pipeline.

For this purpose we consider a temporal train-test loop where we train the novelty detection models using observations within a fixed window size of 2 months and we test their performance on the observations belonging to the next month. Then, the window is displaced using a step size of 1 month, the models are retrained and retested with the new data. This process is repeated for the whole available time period, resulting in 22 train-test iterations.

Regarding the training data and depending on the results obtained from the clustering analysis, we simulate a real-world scenario where we train only with samples from a selected set of clusters, to check if the novelty detection models are capable of detecting the novel clusters over time. In every train step we consider an outlier contamination of 1% during training.

To evaluate the performance of both models we use two metrics: the detection probability or true positive rate (TPR) and the false alarm rate or false positive rate (FPR). The detection probability measures the amount of true novel observations correctly identified by the model while the false alarm rate yields the proportion of common or usual observations that have been wrongly detected as novel.

Considering the ground truth labels to be the labels assigned to each MALDI-TOF MS sample during clustering, a positive prediction is classified as a novel observation and a negative prediction is considered to be typical data. Furthermore, we consider common samples to be those belonging to the training clusters while abnormal observations belong to the remaining groups. Note that we do not make any type of distinction between different novel clusters, we just consider all of them to belong to the same abnormal class, reducing the problem to a simple binary classification.

These metrics are computed for every train-test iteration and averaged at the end to report global values. However, to avoid skewing the mean values we consider two exceptions: for the mean TPR value calculation, only those iterations with outliers in the testset (according to the clustering labels) are considered. Regarding FPR values, train-test iterations with no training cluster samples in the testset are not used in the computation.

Moreover, 95% confidence intervals (CI) are also reported for each performance metric. We use the standard error mean (SEM) method as per Eq. (5) considering a sample size equal to the number of train-test iterations used to compute the mean score value in each case.

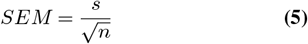

Where *s* is the standard deviation of the sample and *n* is the sample size. Then, 95% CI are obtained as:

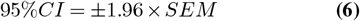

## Results

### K. Preprocessing and clustering

Running the silhouette score evolution analysis on the E-PIKE matrices yields the results represented in Fig. 3. In a general sense, the cohesion of the partitioning decreases rapidly with the number of clusters. The masked version of the pipeline shows the highest scores, followed by the no SPR variant and our proposed method.

**Fig 3.**
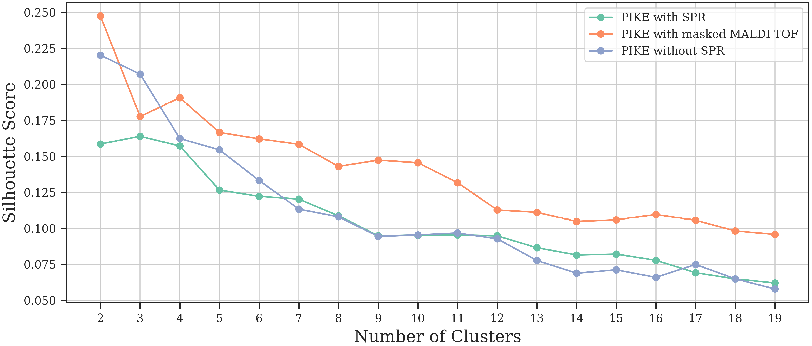
Silhouette score evolution with 20 clusters for all possible kernelization pipelines.

The best score is achieved in both masked peaks and no SPR scenarios when two clusters are used, while the SPR method yields three clusters. However, there is not a significant difference in the silhouette score obtained when using two, three or four groups in this last case.

In light of these results we choose to use 3 clusters in every scenario. Here we prioritize the number of clusters over the silhouette score as differences between strains might be subtle, hence we prefer to overestimate the number of clusters. These clusters have been sent to IiSGM to genomically analyze samples from each group to determine if they are indeed different strains.

Fig. 4 shows the kernel matrices after clustering, ordered per cluster and according to the grouping performed by each pipeline. The SPR pipeline produces a clustering with groups presenting strong intra-cluster cohesion but very similar to each other. However, the third cluster is clearly more isolated than the remaining ones.

**Fig 4.**
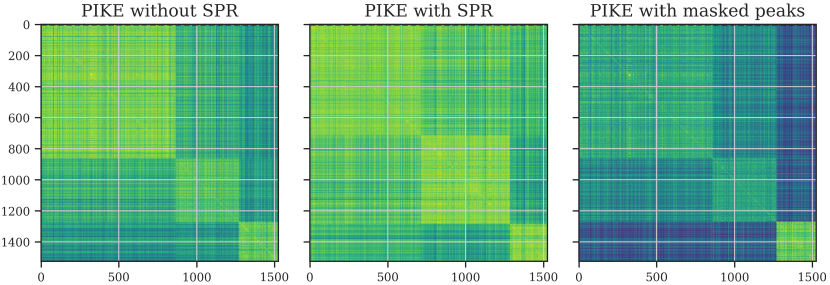
E-PIKE similarity matrices ordered according to cluster assignment.

Regarding the no SPR pipeline, the differences are more prominent. It produces a densely populated and very cohesive base cluster, followed by two smaller and similar in size groups. The second cluster presents similarities with the base group while the third one is more isolated. In this case, the intra-cluster cohesion is as strong as the one yielded by its SPR counterpart while the similarities among clusters are less prominent.

With respect to the masked pipeline, the clustering is very similar to the pipeline without SPR, but with less intra-cluster cohesion while cluster 2 is highly isolated.

Table 2 presents the number of samples per cluster and pipeline. Both no SPR and masked peaks pipelines present a very similar number of samples per cluster while their SPR counterpart shows greater differences in the samples assigned to clusters 0 and 1.

**Table 2.** Number of Samples per Cluster and Pipeline.

|  | Cluster 0 | Cluster 1 | Cluster 2 |
| --- | --- | --- | --- |
| **SPR** | 715 | 569 | 241 |
| **No SPR** | 864 | 407 | 254 |
| **Masked Peaks** | 858 | 412 | 255 |

Performing a more detailed analysis of the clustering reveals that 233 samples are always assigned to cluster 0, only 1 sample is consistent in cluster 1 for all three pipelines and 232 samples are always present in cluster 2.

Fig 5 presents the mean MALDI-TOF spectra, per cluster, for these consistent samples across pipelines.

**Fig 5.**
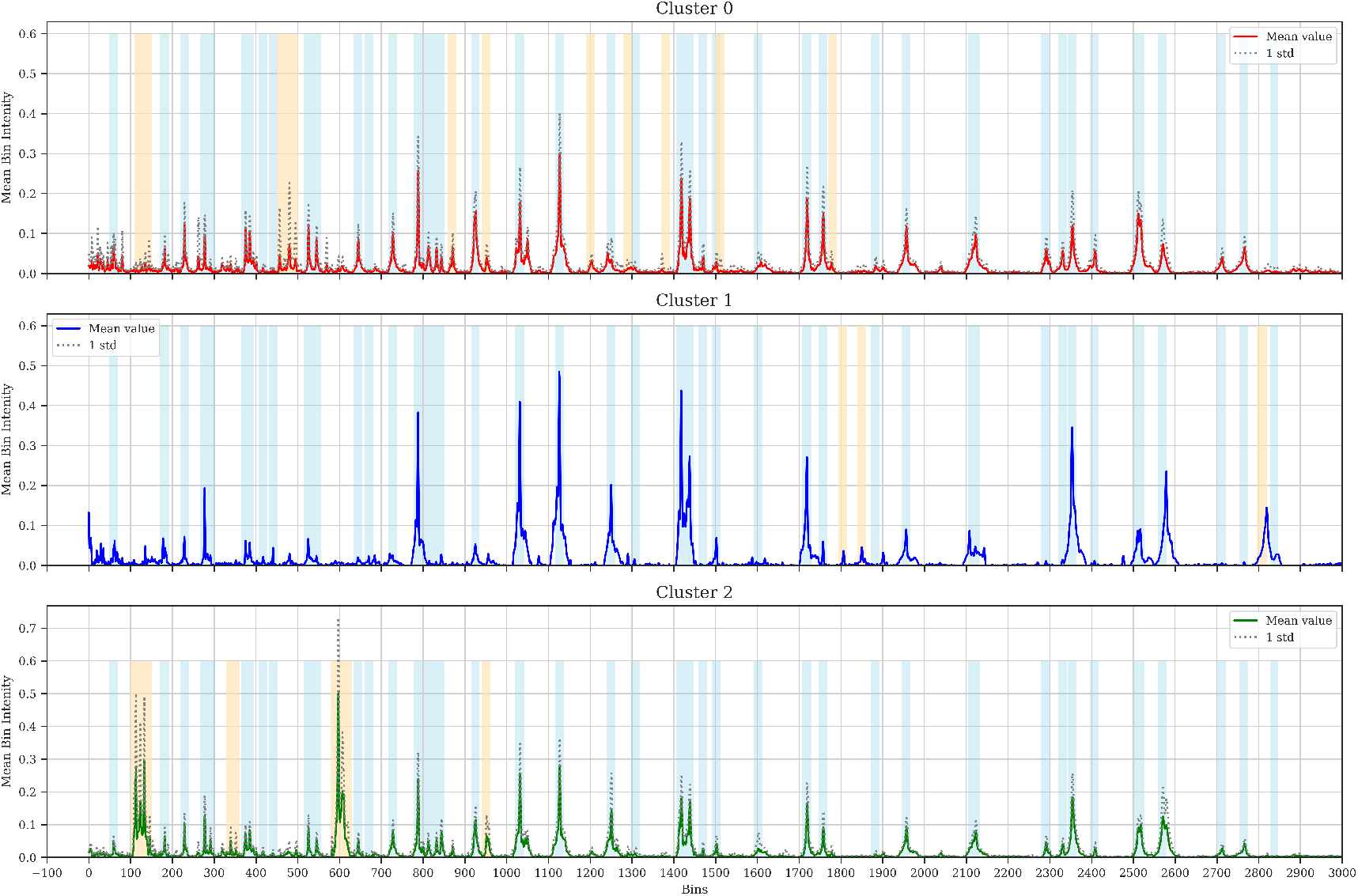
Mean MALDI-TOF spectra, per cluster, of the samples that are always assigned to the same cluster. Blue shaded regions represent common peaks identified by IiSGM. Orange shaded regions mark non-common peaks that differ between spectra. X-axis ranges in [0, 3000] bins for visualization purposes.

Apart from the variations that common peaks might present, there is a clear difference between samples from the first two clusters and the third one, peaks in the range [100, 150] and around bin 600 are only present with high intensity in the last group. In the other hand, the differences between clusters 0 and 1 are more subtle and are associated with less relevant peaks, as shown by the orange shaded areas in Fig 5.

Fig. 6, Fig. 7 and Fig. 8 show the mean MALDI-TOF spectra of the non-intersecting samples, per cluster, considering the partition done by our SPR proposal, the E-PIKE without SPR and the E-PIKE with masked common peaks, respectively.

**Fig 6.**
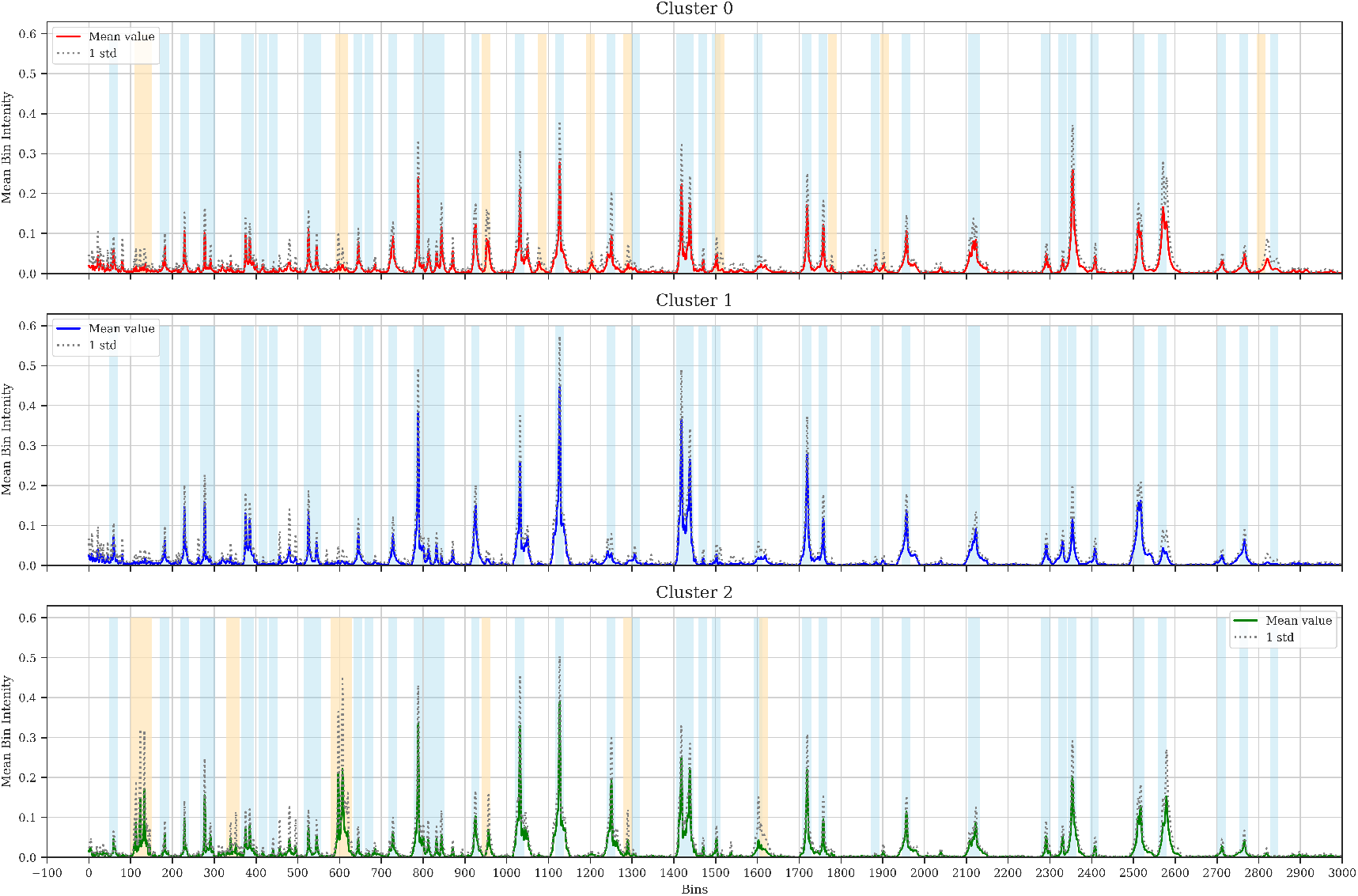
Mean MALDI-TOF spectra, per cluster, of the non-intersecting sample partition by our SPR proposal. Blue shaded regions represent common peaks identified by IiSGM. Orange shaded regions mark non-common peaks that differ between spectra. X-axis ranges in [0, 3000] bins for visualization purposes.

**Fig 7.**
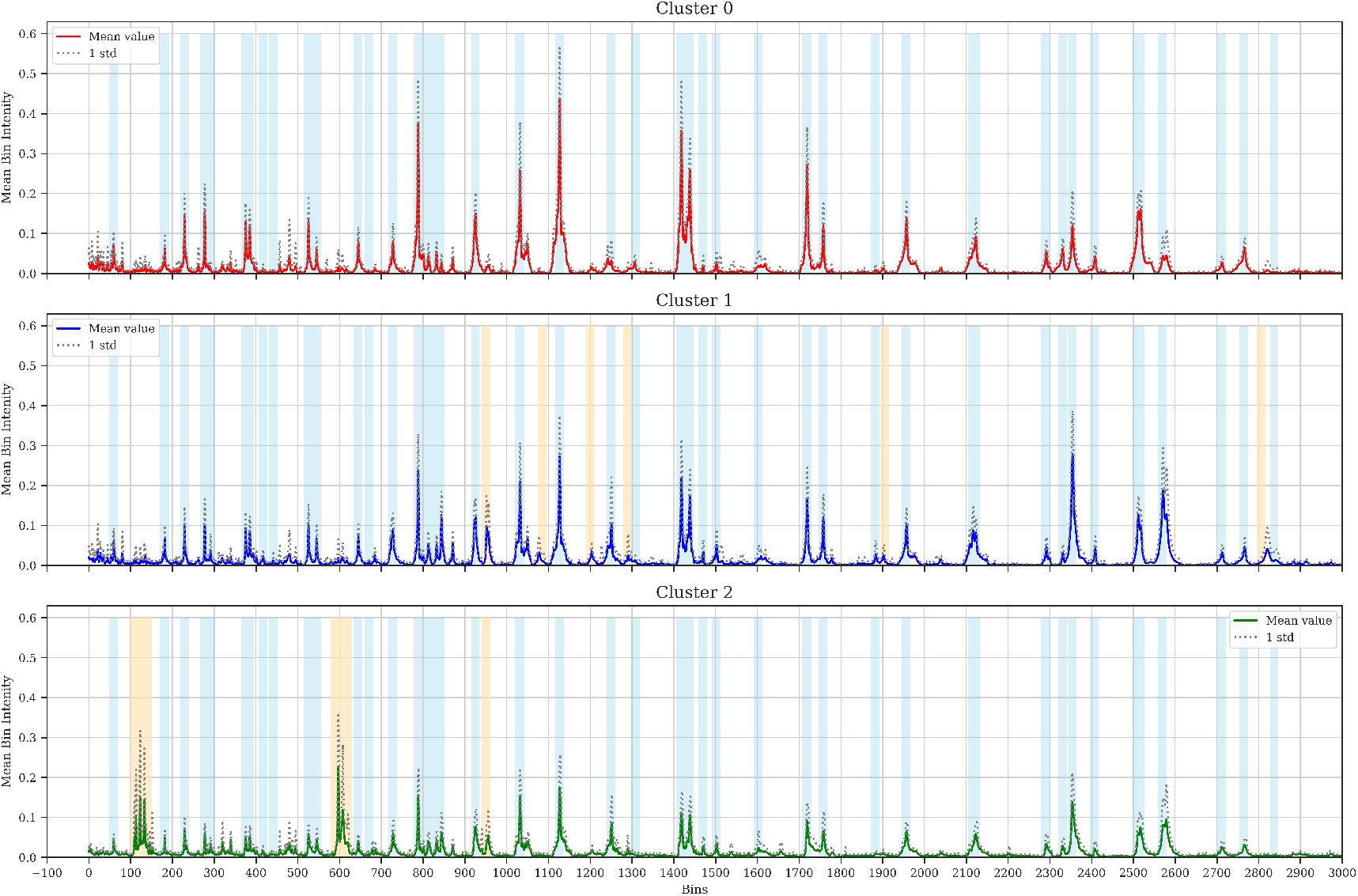
Mean MALDI-TOF spectra, per cluster, of the non-intersecting sample partition by the no SPR pipeline. Blue shaded regions represent common peaks identified by IiSGM. Orange shaded regions mark non-common peaks that differ between spectra. X-axis ranges in [0, 3000] bins for visualization purposes.

**Fig 8.**
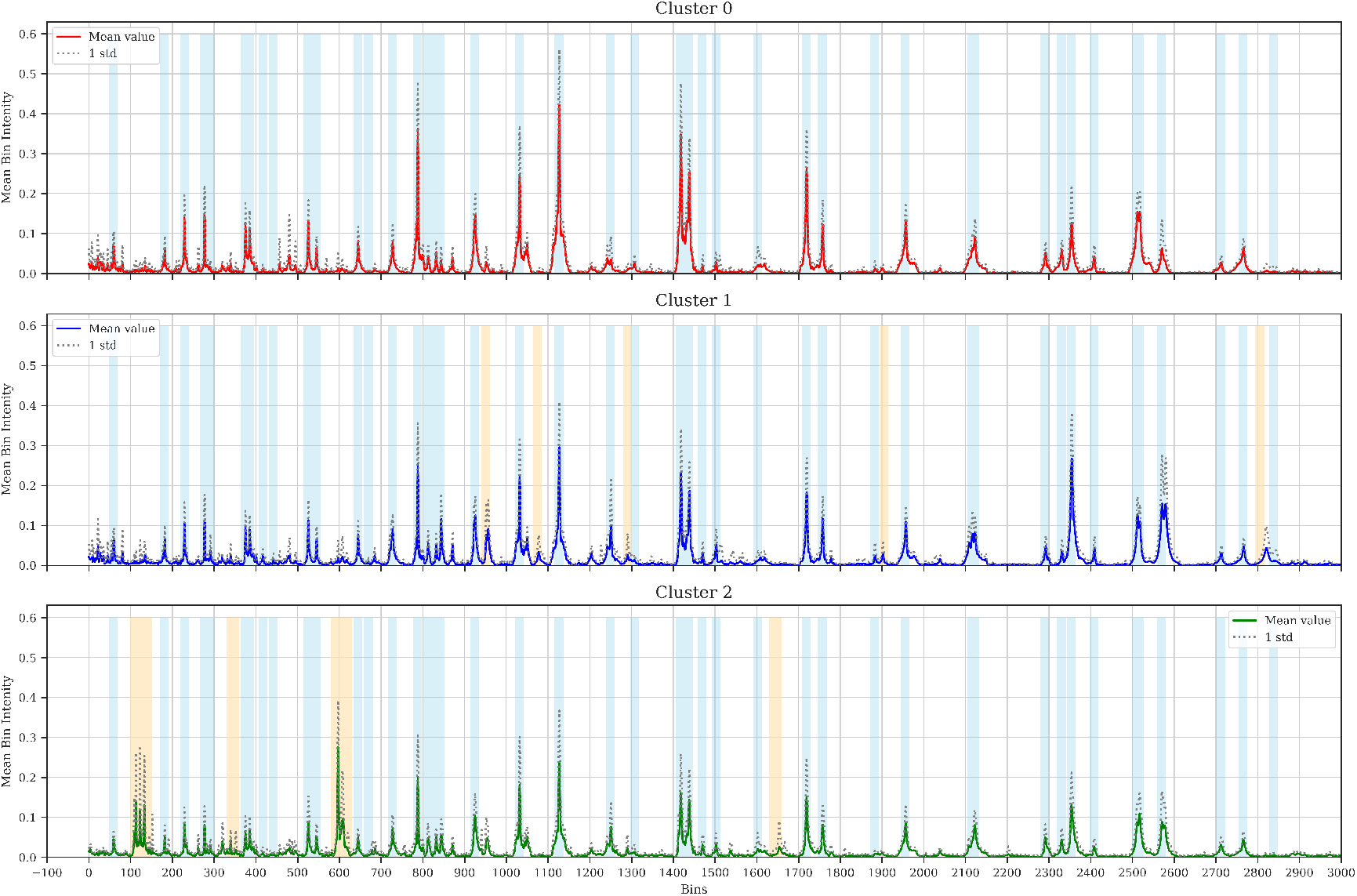
Mean MALDI-TOF spectra, per cluster, of the non-intersecting sample partition by the masked pipeline. Blue shaded regions represent common peaks identified by IiSGM. Orange shaded regions mark non-common peaks that differ between spectra. X-axis ranges in [0, 3000] bins for visualization purposes.

Focusing on cluster 2, it is possible to appreciate that the same high intensity peaks identified in the intersecting samples are present in all three pipelines. The difference in group assignment between pipelines is caused by secondary peaks. The SPR pipeline is able to highlight more peaks outside the common peak regions, as shown by the shaded orange regions.

This same trend is present in the way the different pipelines partition the non-intersecting samples for clusters 0 and 1.Our proposed method presents higher inter-cluster differences in the non-common peaks while the two remaining pipelines tend to yield more similar spectra, with non-common peaks being more similar.

It is possible to identify this behaviour, for example, in the peaks that appear around bin 600 for clusters 0 and 1, our SPR proposal shows higher peak intensities for cluster 0 while the other two pipelines present very similar peak height. Another example of this same phenomenon can be seen around bin 100, again for the first two clusters.

This results show that the combination of an E-PIKE and SPR method is better at highlighting the differences in the non-common peak regions.

### L. Novel strain identification

In light of the previous results we choose the E-PIKE with SPR to test the OCSVM and iForest models. We train the models using only samples from cluster 0 and considering as novel samples from clusters 1 and 2. Even though the differences between clusters 0 and 1 are not so prominent we choose this training scheme to consider the most general case possible.

Fig. 9 summarizes the results obtained in terms of detection probability and false alarm ratio. The OCSVM model achieves a clearly higher TPR score, being more effective and consistent at detecting novel strains. However, regarding the FPR score, the iForest outperforms the OCSVM with a much lower proportion of non-novel samples identified as novel.

**Fig 9.**
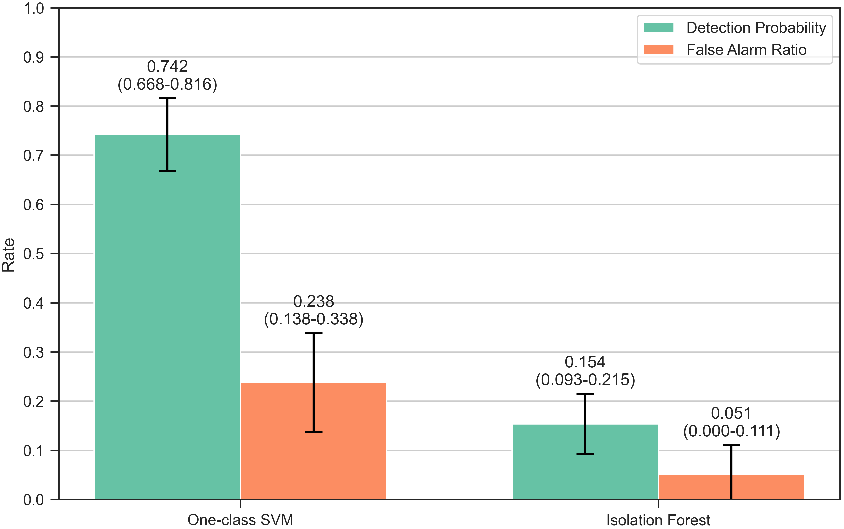
Mean detection probability and false alarm ratio scores, for every train-test iteration, obtained by the novelty detection models. 95% confidence intervals are included in parentheses.

## Discussion

The results revealed several key findings. First, we found that our proposed E-PIKE with SPR in kernel space is able to highlight peak position and intensity differences in the non-standard *E. coli* peak regions, while the remaining baseline pipelines present more similar peaks within these regions of interest; second, we proved that at least one cluster presents almost the same samples in every pipeline and that its mean MALDI-TOF spectrum shows novel proteins with respect to the other two groups in the [100, 150] and [580, 620] bin regions, being a strong strain candidate. This findings validate that the combination of, at least, our

E-PIKE implementation and spectral clustering techniques is sufficient enough to discern isolates below the species level when the differences are prominent. Finally, we confirmed that a OCSVM novelty detection model, in combination with the E-PIKE and SPR method, is able to identify novel isolates in an unsupervised manner with a high rate of detection probability even when the differences between clusters are not outstanding.

The observed differences between clustering pipelines, mainly related to clusters 0 and 1 can be explained attending to the basic principle of SPR. The main objective is to remove common information within the spectra to point out the differences in the non-common peak regions where the subspecies information is contained. Thus, maintaining the highest peaks will cause the kernel to assign similarities based, mainly, on the intensity differences these peaks might present.

Following this analysis one might argue that this same behaviour should be identified in the kernel matrices, finding more sparsity in the SPR pipeline, however our results show the opposite behaviour. When removing peaks within PIKE space, the only differences between spectra are due to small and very spare peaks, contrary to the big differences that can be found between common peaks. These superior differences are mainly caused by the intensity variability in common peaks inherent to MALDI-TOF data.

Regarding the masked pipeline, in theory it might seem to be a good approach. However, as we have proven, it does not show very different results to the ones generated by the no SPR pipeline. The main cause for this phenomenon is that the localization and width of the common peaks is not deterministic but might vary slightly between spectra, reducing the accuracy of the masking.

This problem is not such when removing peaks in PIKE space as the very basic formulation of the kernel takes into account these slight position and width differences by means of its smoothing hyperparameter *t*. Yet, the lack of ground-truth labels obtained from a golden standard method did not allow us to perform a proper cross-validation of this parameter to maximize peak differences. We leave this proposal for a future project.

Altogether, these results prove the superiority of our proposed new methodology, the combination of an E-PIKE and the SPR method. It is common practice in the literature to only work with a subset of the highest peaks. However we can conclude, based on our results, that it is better to remove common peaks in kernel space to highlight non-common peak differences and, consequently, the subspecies information.

We have also found a strong *E. coli* strain candidate, cluster 2 is formed, in all three pipelines, by almost the same samples while the partition of clusters 0 and 1 depends on the pipeline. Furthermore, the mean MALDI-TOF spectrum of cluster 2 shows significant differences with the other groups in the [100, 150] and [580, 620] bin regions.

However, these results are not yet confirmed by IiSGM, which is genetically sequencing stored bacteriological isolates from each cluster to confirm whether the groups we found correspond to different strains or the differences are provoked by other MALDI-TOF MS sources of variability. In a future study, once we have obtained the gold standard labels and the number of strains contained within the data, we will be able to perform a more accurate clustering and a thorough analysis of the strains to identify possible candidate peaks for *E. coli* strain identification.

Regarding novel strain identification, we have shown that a OCSVM combined with the E-PIKE implementation and the SPR method is able to detect novel groups not present in the training data in an unsupervised manner, even when the differences between these strains are not significant. The model is able to detect novel samples from clusters 1 and 2 with a high degree of TPR.

In this regard, our proposed method has the potential to facilitate the process of deciding which samples to sequence. Genetic sequencing of bacteria is costly, hindering the routine testing of all bacteriological isolates acquired during standard clinical practice. Therefore, our combination of an E-PIKE, SPR and a OCSVM could be employed to detect unusual isolates and, subsequently, sequence only those identified as rare. This approach allows for targeted sequencing, focusing on samples flagged as anomalous rather than performing blind sequencing.

The difference in TPR performance between the OCSVM and the iForest models can be explained by the underlying mechanisms of both models. One-class SVMs are inherently compatible with kernel methods while iForests are not designed to support this compatibility. Nonetheless, the iForest presents a better performance in terms of FPR, being more effective in correctly classifying non-novel samples as such.

Note that these results in terms of difference in model performance are somehow predictable since the typical/novel labelling has been made with the clustering results of the E-PIKE and SPR pipeline. We will be able to confidently validate these results when the golden standard labels are available.

Furthermore, the hyperparameters of both models have not been cross-validated in any way but assigned common values that usually work well. The lack of a priori and reliable ground truth labels has hampered the optimization of these parameters, influencing the performance of both models, which might have caused the underperformance of the OCSVM in the FPR score. However, the findings shown in this study suggest a promising line of work.

## Conclusions

We have proposed, to our knowledge, the first method for unsupervised novel bacterial identification specifically tailored to strain identification, showing promising results. In addition we have demonstrated the ability of our system to detect novel bacteriological isolates within clinical settings using MALDI-TOF MS data in combination with a customized E-PIKE implementation, our newly proposed SPR method, clustering techniques and novelty detection models.

Consequently, our approach allows for the optimization of the genetic sequencing process during standard clinical practice. By only focusing on rare and anomalous samples it is possible to perform a targeted sequencing, reducing costs and facilitating the clinicians’ decision making process.

First, we have introduced E-PIKE, an efficient modification of the original PIKE implementation. This implementation allows for the computation of kernel matrices using complete MALDI-TOF spectra, contrary to the standard practice in the literature that uses a subset of the most relevant peaks. Specifically, the original authors use a subset of the200 most relevant peaks, while our implementation is able to compute affinity matrices with 6000 dimensional spectra, without loosing computational tractability and incrementing by a factor of 30 the capacity of the kernel.

Second, we extended the PIKE calculation to adapt it to the problem of strain identification by introducing the SPR step. Then, we demonstrated its ability to highlight small differences at the subspecies level in the less relevant peak regions, proposing three different *E. coli* groups, with one of them being a strong strain candidate.

Finally, we proved the ability of a OCVSM model, in combination with our E-PIKE implementation and the SPR adaptation, to detect rare isolates with a high rate of detection probability.

This findings have the potential to leverage the power of ML to detect HAIs the very first moment they happen to take the necessary measures to isolate them at the appropriate time.

## ACKNOWLEDGEMENTS

Not yet.

